# Resting-state connectivity underlying cognitive control’s association with perspective taking in callous-unemotional traits

**DOI:** 10.1101/2022.03.24.485718

**Authors:** Drew E. Winters, Daniel R. Leopold, R. McKell Carter, Joseph T. Sakai

**Affiliations:** Department of Psychiatry, University of Colorado School of Medicine, Anschutz Medical Campus; Department of Psychology & Neuroscience, University of Colorado Boulder, Boulder, CO, USA; Institute of Cognitive Science, University of Colorado Boulder, Boulder, CO, USA; Department of Electrical, Computer and Energy Engineering, University of Colorado Boulder, Boulder, CO, USA

**Keywords:** Cognitive Control, Perspective Taking, Callous-Unemotional Traits, GIMME, Functional Connectivity

## Abstract

Callous-Unemotional (CU) traits are often associated with impairments in perspective taking and cognitive control (regulating goal directed behavior); and adolescents with CU traits demonstrate aberrant brain activation/connectivity in areas underlying these processes. Together cognitive control and perspective taking are thought to link mechanistically to explain CU traits. Because increased cognitive control demands modulate perspective taking ability among both typically developing samples and individuals with elevated CU traits, understanding the neurophysiological substrates of these constructs could inform efforts to alleviate societal costs of antisocial behavior. The present study uses GIMME to examine the heterogenous functional brain properties (i.e., connection density, node centrality) underlying cognitive control’s influence on perspective taking among adolescents on a CU trait continuum. Results reveal that cognitive control had a negative indirect association with CU traits via perspective taking; and brain connectivity indirectly associated with lower CU traits – specifically the social network via perspective taking and conflict network via cognitive control. Additionally, less negative connection density between the social and conflict networks was directly associated with higher CU traits. Our results support the growing literature on cognitive control’s influence on socio-cognitive functioning in CU traits and extends that work by identifying underlying functional brain properties.

## 1. Introduction

Callous-unemotional (CU) traits are a youth antisocial phenotype associated with persistent violent and criminal (i.e., antisocial) behavior (Blair et al., 2014) that relate to affective (i.e., emotional) deficits of adult psychopathy involving poor affective response and empathy for others (Barry et al., 2000). Central CU traits impairments involve perspective taking (i.e., understanding another by adopting their perspective [Davis, 1980; 1983]; Anastassiou-Hadjicharalambous & Warden, 2008; Lui et al., 2016; O’Kearney et al., 2017) and cognitive control (also called executive control; Baskin-Sommers et al., 2015; Gluckman et al., 2016; Winters & Sakai, 2021). Cognitive control impairments relate to antisociality (Zeier et al., 2012) and is necessary to modulate socio-cognitive processes (for reviews see: Mahy et al., 2014; Wade et al., 2018) such as perspective taking (Lamm et al., 2010; Qureshi & Monk, 2018). Decrements in perspective taking associated with CU traits is exacerbated under increased cognitive control demands (Winters & Sakai, 2021). These impairments appear to partially result from differences observed in brain regions associated with perspective taking (e.g., Thijssen & Kiehl, 2017; Umbach & Tottenham, 2020) and cognitive control (e.g., Pu et al., 2017; Szabó et al., 2017; Yoder et al., 2016), but these neuroimaging results fail to adequately account for parametric brain pattern heterogeneity in youth with CU traits (e.g., Winters, Sakai, et al., 2021). Overall, cognitive control impairments robustly associate with antisocial behavior, and CU traits are a developmentally useful and clinically meaningful precursor to antisocial behavior. Given the inherent harm to others and society, it is imperative to fully understand what aspects of behavior and biology might explain this association and contribute to later violent and criminal behavior. While much has been learned about the neural underpinnings of antisocial and delinquent behavior among adults, the need for generalizing these findings to adolescents, accounting for individual heterogeneity of brain patterns (e.g., Winters, Sakai, et al., 2021), and pinpointing malleable intervention targets remains (Herpers et al., 2014). As such, the present study examines functional properties of adolescent brains and perspective taking – an important and impoverished ability among adolescents with elevated CU (Lui et al., 2016) – as a plausible explanation for the association between CU traits and cognitive control.

### 1.1. Cognitive Control

Cognitive control is differentiated from the umbrella of executive functions in its intentional relation toward goal directed behavior as opposed to habitual cognitive processes (Friedman & Robbins, 2022) that involve monitoring conflict to gage the demand for control and then using information from the present context to regulate goal directed behaviors (Botvinick et al., 2001). Impairments in cognitive control, and damage to the anterior cingulate cortex (ACC) in particular, reduces the capacity of resolving conflict necessary to respond to other’s emotions and subsequent response to other’s emotions (Maier & di Pellegrino, 2012). A lack of response to others’ emotions is considered a fundamental pathway for psychopathy (Blair, 2008; Blair & Mitchell, 2009). Interpreting this lack of emotional response has largely centered on the existence of profound affective impairments (Blair, 2008; Blair & Mitchell, 2009), but there is substantial evidence it can be explained by a “bottleneck” in the early stages of attention processing (i.e., cognitive control; Braver, 2012) blocking peripheral information processing (i.e., the response modulation hypothesis; Lorenz & Newman, 2002).This effect has been demonstrated using conflict paradigms measuring cognitive control, such as the Stroop or Flanker task, that are modified to include a spatial component for peripheral stimuli (e.g., Gluckman et al., 2016; Vitale et al., 2005). These studies suggest that CU traits are associated with decrements in cognitive control (Gluckman et al., 2016) that are especially prominent during high conflict, incongruent task conditions (Botvinick et al., 2001; Yeung, 2013).

Brain regions recruited during conflict paradigms measuring cognitive control center around the anterior cingulate cortex (ACC) and pre-supplementary motor area (pre-SMA) forming a conflict network. The ACC is a central region for detecting conflicts in information processing and signaling top-down control (e.g., Brown, 2013; Shenhav et al., 2013; Yeung, 2013). The pre-SMA is thought to play a leading role in response-based conflict (Egner et al., 2007; Nachev et al., 2007) by evaluating outcomes of actions (Bonini et al., 2014). The ACC and pre-SMA are particularly sensitive to conflict during the Stroop task (Banich, 2019; Milham & Banich, 2005) and have been used to evaluate conflict sensitivity (e.g., Yang et al., 2021). These specific ROIs are used to investigate cognitive control during flanker tasks with spatial stimuli (e.g., Iannaccone, Hauser, Ball, et al., 2015; Iannaccone, Hauser, Staempfli, et al., 2015). The dorsal medial prefrontal cortex has also been implicated (e.g., Alexander & Brown, 2011). However, others have argued that this activation reflects a longer task duration rather than a conflict response (Grinband et al., 2011a, 2011b), which has been supported empirically using both EEG and fMRI (Iannaccone, Hauser, Staempfli, et al., 2015). As such, many studies have not included the dorsal medial prefrontal cortex when investigating conflict paradigms to assess cognitive control (e.g., Iannaccone, Hauser, Ball, et al., 2015; Iannaccone, Hauser, Staempfli, et al., 2015).

Functional abnormalities in the ACC and pre-SMA among youth with CU traits further supports the presence of cognitive control impairments. For example, the ACC demonstrates less activity during facial emotion recognition tasks (Szabó et al., 2017) and pain responses (Marsh et al., 2013) as well as less functional connectivity seeded in the ACC (Yoder et al., 2016). Similarly, the pre-SMA demonstrates decreased activation when responding to others’ emotions (O’Nions et al., 2017) and viewing another person in pain (Decety et al., 2013), as well as aberrant functional connectivity of the pre-SMA (Pu et al., 2017) at higher levels of CU traits. It is therefore pertinent to focus on the ACC and pre-SMA to understand cognitive control in relation to CU traits.

Defining neurophysiological substrates of cognitive controls influence on perspective taking in relation to CU traits is critical for understanding mechanisms of antisocial behavior. Given that the above literature suggests functional coupling between the ACC and pre-SMA supports cognitive control during conflict paradigms, we hypothesize that those with elevated CU traits would have disrupted connectivity whereas greater connectivity would positively associate with cognitive control and indirectly impact CU traits. Such evidence could reveal brain function underlying cognitive control impairments associated with CU traits; and this investigation could be further extended by accounting for perspective taking.

### 1.2. Perspective taking

Perspective taking, or capacity to adopt another’s point of view and attributing their thoughts and feelings (Decety, 2011), is supported by distinct brain regions and is critical for healthy social (Decety, 2005), moral (Decety & Cowell, 2014), and prosocial behavior (Decety et al., 2016; Tamnes et al., 2018). CU trait impairments in perspective taking (Anastassiou-Hadjicharalambous & Warden, 2008; Lui et al., 2016; O’Kearney et al., 2017) predict antisocial behavior above clinical ratings of CU traits (Gillespie et al., 2018; Song et al., 2016). CU trait impairments in perspective taking have been observed in the brain. For example, brain regions supporting perspective taking consist of the temporal parietal junction (TPJ), medial prefrontal cortex (mPFC) and the posterior cingulate cortex (PCC) (for meta-analysis: Fehlbaum et al., 2021) that form a social network (Blakemore, 2012; Klapwijk et al., 2013; McCormick et al., 2018); and, youth with elevated CU traits demonstrate aberrantly decreased functional connectivity among these regions (Thijssen & Kiehl, 2017; Umbach & Tottenham, 2020; Winters & Hyde, 2022). Although underlying multiple cognitive functions, these regions’ associations with perspective taking are thought to reflect cognitive processes of reflection and understanding mental states of oneself and others’ (Buckner et al., 2008; Buckner & Carroll, 2007; Uddin et al., 2009). Less connectivity in this network may indicate difficulty inferring others’ cognitive and emotional states; whereas greater connectivity in the social network is associated with greater perspective taking in adolescents (Winters, Pruitt, et al., 2021). Thus, greater connectivity in the social network could be associated with lower CU traits indirectly via perspective taking. Such evidence could reveal brain patterns underlying social processing impairments associated with CU traits. Moreover, core decrements associated with CU traits could be revealed by examining these brain association with cognitive control.

### 1.3. Cognitive Control and Perspective Taking

Substantial evidence supports that perspective taking is supported by cognitive control in typically developing samples. For example, those with higher inhibitory control report higher perspective taking (Carlson, 2005; Lamm et al., 2010; Wade et al., 2018); and selecting between different stimuli to perform perspective taking is reliant on cognitive control (Qureshi & Monk, 2018; Qureshi, Monk, et al., 2020). Numerous studies converge on these links suggesting that cognitive control modulates perspective taking (for reviews see: Mahy et al., 2014; Wade et al., 2018). Perspective taking is thought to be modulated by (1) processing the emotional context and (2) resolving conflicts between one’s own and others emotions (Deschrijver & Palmer, 2020).

Neuroimaging studies support links between cognitive control and perspective taking. Processing conflict during perspective taking has shown causal associations in transcranial magnetic stimulation of the dorsolateral prefrontal cortex (dlPFC) in typical samples (Kalbe et al., 2010; Qureshi, Bretherton, et al., 2020) and in those with CU traits (Konikkou et al., 2020), which can be influenced by exerting cognitive control (e.g., Seymour et al., 2018). Thus, it is plausible that social and conflict network connectivity with the dlPFC may underlie cognitive control and perspective taking impairments amongst youth with CU traits.

Youth with CU traits demonstrate connections between social and conflict networks that may account for impairments in the cognitive control and perspective taking link. In typically developing samples we expect anticorrelation between the social and conflict networks because regions of the conflict network are active when engaged in a task (i.e., task-positive network) whereas social network regions are active when not engaged in an external task (i.e., task-negative network; Uddin et al., 2009). However, youth with CU traits demonstrate less anticorrelation between task-positive and task-negative networks (Pu et al., 2017; Winters, Sakai, et al., 2021), which may represent developmental immaturity (Richardson et al., 2018). Given the importance of cognitive control for perspective taking, it is plausible that functional coupling observed with greater anticorrelation between these networks may be important for cognitive control and perspective taking impairments in CU traits.

Understanding neurophysiological underpinnings of how cognitive control and perspective taking are linked to CU traits would have tremendous value in the public health domain. We previously tested how affective theory of mind (a socio-cognitive process related to perspective taking) is affected when placing additional demands are placed on cognitive control. This study revealed that those higher in CU traits had greater difficulty inferring others’ emotions after taxing cognitive control (Winters & Sakai, 2021). Our study demonstrates a vulnerability in cognitive control that impacts theory of mind accuracy; however, it does not answer whether this vulnerability is because of cognitive control and perspective taking directly interacting nor does it identify the neurophysiological underpinnings of this relationship. Further investigation into these unanswered questions can help clarify the core behavioral and neural processes underlying CU traits, which may in turn provide valuable predictive and intervention-related information.

### 1.4. Methodological Improvements for Estimating Functional Brain Properties

Previous studies characterizing the brain in relation to constructs of interest could be improved by modeling heterogeneity of individual connectivity in adolescent brains. For example, functional brain patterns are as unique as fingerprints (Damoiseaux et al., 2021). Ignoring this heterogeneity when modeling brain features can cause spurious associations, whereas modelling the heterogeneity of individual brains can more accurately characterize network patterns (Gates & Molenaar, 2012). Additionally, lagged connections, or the temporal relationship such that a prior timepoint of one brain node predicts the future timepoint of another brain node, are often overlooked in traditional approaches but are an important feature characterizing brain function (Mitra et al., 2014) and help improve estimation of contemporaneous connections. Thus, the present study uses a method that accounts for heterogeneity and both lagged and contemporaneous connections called subgrouping group iterative multiple model estimation (S-GIMME; Gates et al., 2017). S-GIMME and GIMME outperform other network modeling approaches (e.g., Bayes nets and Granger causality; Gates et al., 2017; Gates & Molenaar, 2012), but here we use S-GIMME because it improves estimation of individual level connections over GIMME by providing the model with more known priors (Beltz & Gates, 2017). These methods have been used to demonstrate brain heterogeneity in psychopathy (Dotterer et al., 2020) as well as CU traits (Winters, Sakai, et al., 2021). Thus, it is appropriate to leverage S-GIMME to characterize functional brain properties of CU traits and underlying psychological processes supporting these symptoms.

In addition, prior work investigating the brain and CU traits primarily relies on task-based activations and our use of S-GIMME could meaningfully build on this prior work by incorporating a contemporary understanding of brain function. As opposed to the modular view of the brain espoused by task-based studies making up the majority of investigations on CU traits and the brain, contemporary views on the brain recognize that human cognition requires the integration of multiple brain regions (McIntosh, 2000), forming networks supporting brain function and that understanding the functional connectivity across these regions and networks can provide a deeper insight into human behavior (Bassett & Sporns, 2017). This functional connectivity represents the distributed function among brain regions (Zhang et al., 2021), that capture critical developmental features of adolescent brains (Ernst et al., 2015; Uddin et al., 2011). Investigating adolescent brains using S-GIMME will quantitate individual level functional connectivity while allowing us to derive network properties of node centrality and network density. Network density quantifies the number of connections that exist between the set of nodes making up a network with higher values indicating more paths where information can be transferred between nodes of that network and lower values indicating fewer paths for transferring information between nodes. The density of these paths can represent positive or negative connections. Higher positive density indicates regions activating together whereas higher negative density indicates regions activating opposite to each other, or more functional coupling. Node centrality measures the number of connections a node has relative to and potential number of connections it could have, which represents the importance of that node. Positive node centrality indicates the number of positive connections and negative node centrality indicated the number of negative connections.

### 1.5. The Present Study

The present study aims to 1) examine the extent to which cognitive control’s association with perspective taking accounts for CU traits and 2) characterize the related functional brain properties in a community sample of early-to-mid adolescents (ages 13-17). We took an iterative approach by first examining the behavioral model, then the associated functional brain properties, before constructing a final model. To characterize the functional brain properties, we used S-GIMME to generate person specific connectivity maps to derive network density and node centrality for all participants. We hypothesized that cognitive control’s positive association with perspective taking would partially account for CU traits. For functional brain properties, we hypothesized the following: a) elevated CU traits would associate with less positive connection density in the conflict and social networks; b) cognitive control would be positively associated with connection density in the conflict network; and c) perspective taking would positively associate with positive connection density in the social network. For the dlPFC association with social and conflict networks, we hypothesized that negative connections would correlate positively with both perspective taking and cognitive control. For node centrality, the right TPJ is a central node for perspective taking (Martin et al., 2020), so we hypothesized that perspective taking would associate with node centrality in the social network; and, given the ACC is central for detecting conflict, we hypothesized that cognitive control would associate with centrality in the ACC. Finally, we anticipated that less negative density between the social and conflict networks would associate with CU traits and greater negative density would associate with cognitive control and perspective taking.

## 2. Methods

### 2.1. Sample

Participants were sampled from the Nathan Kline Institute’s Rockland study downloaded from the 1000 connectomes project (www.nitrc.org/projects/fcon_1000/). We included all participants between the ages of 13-17 with a WAIS-II (α = .96; Wechsler, 2011) IQ score of >80. From a total of 122 participants, we removed 10 for IQ < 80 leaving 112 participants for analysis. Youth in the sample were predominantly White (White= 63%, Black = 24%, Asian = 9%, Indian = 1%, other= 3%) with marginally more males (female = 43%) and an average age of 14.52±1.31 years. Written consent and assent was obtain from all participants (for study procedures see: Nooner et al., 2012).

### 2.2. Measures

#### Interpersonal Reactivity Index (IRI)

Perspective taking was assessed using the IRI perspective taking subscale (Davis, 1980, 1983). Perspective taking is defined as the tendency to adopt others psychological point of view (e.g., “I try to look at everybody’s side of a disagreement before I make a decision”; present sample α=.74). Higher scores indicate higher dispositional perspective taking.

#### Inventory of Callous-Unemotional Traits (ICU)

The total score of the 24-item ICU was used to assess CU traits (Frick, 2004). We used the same factor structure that was validated by Kimonis et al. (2008) that removed two items due to poor psychometrics. This factor structure had an adequate reliability in the current sample (present sample α=.72). Higher scores indicate greater CU traits.

#### Attention Network Task (ANT)

Cognitive control was measured using the ANT task by an index of conflict between incongruent versus congruent flanker conditions (Fan et al., 2002; Rueda et al., 2004). Participants were presented with five arrows either above or below a central fixation cross and are asked to determine the direction of the target arrow that is presented in the middle – either left or right. The task had two flanker conditions that were congruent or incongruent indicating whether the target arrow was consistent with the initial arrows or not. After a fixation window, a cue was presented for 300ms. Then, after a variable duration (300 – 6,300ms), the stimulus appears either above or below the fixation point where the participant has up to 2,000ms to respond with the direct of the target arrow. The inter trial interval was also variable (from 3,000 – 15,000ms). There were 72 trials for congruent and incongruent conditions resulting in a total of 144 trials.

Covariates are described in Supplemental Methods. Importantly, conduct problems were included as a covariate (see Supplemental Methods).

### 2.3. Imaging measures

Methodological information on imaging acquisition, preprocessing, and S-GIMME are described in Supplementary Methods.

#### Region of Interest Selection

We selected regions corresponding to the social and conflict networks. For the social network, we used predefined ROIs found across studies to be involved in perspective taking (for meta-analysis: Fehlbaum et al., 2021) consisting of the TPJ, mPFC, and the PCC that are central components of the social brain network (Blakemore, 2012; Burnett et al., 2009; Burnett & Blakemore, 2009; Klapwijk et al., 2013; McCormick et al., 2018). For the conflict network, we used the predefined ROIs involved in a conflict paradigm that measures cognitive control in adolescents, which included the ACC and pre-SMA (Iannaccone, Hauser, Ball, et al., 2015; Iannaccone, Hauser, Staempfli, et al., 2015). The dlFPC was also included because network communication with the dlPFC may be causally related to resolution of conflict in perspective taking (Kalbe et al., 2010; Qureshi, Bretherton, et al., 2020); thus was not included in the definition of either network but was used to probe the network associations with dlPFC.

### 2.4. Analysis

All inferential statistics were conducted in the statistical language R (Version 4.04; R Core Team, 2021) using the ‘lavaan’ (Rosseel, 2012) and ‘GIMME’ (Lane et al., 2021) packages. First, we extracted network features from the S-GIMME networks, then conducted missing data analyses before iteratively building the model using path analyses. Path analyses were estimated using maximum likelihood with Huber-White robust standard errors to correct asymptotic standard errors and improve confidence interval estimation (Maas & Hox, 2004). We estimated bias-corrected bootstrapped parameters with 5000 resamples for all estimated parameters and indirect effects. Importantly, all p-values were bootstrap corrected as a less conservative approach reducing type I error rate without inflating type II errors. We also examined sex as a potential moderator using multigroup models. Finally, we examined node centrality and subgroup associations with CU traits, perspective taking, and cognitive control.

#### Network features

We extracted positive and negative network density as well as node centrality within and between all networks of interest. Network density involved the number of contemporaneous and lagged connections between nodes that were calculated separately for within and between networks. Node centrality was calculated by the number of connections a node has relative to the number of potential connections within a network or between networks (connections / nodes – 1).

#### Missing Data Analysis

Details on assessment and treatment of missing data are described in Supplementary Methods.

#### Model building

Model building took an iterative approach to inform the final full model. At each step we ran a set of likelihood ratio tests for direct and indirect effects to obtain the most parsimonious model while retaining the parameters that best reproduced the data. Prior to all formal model building we outlined hypothesized associations to be tested and ran correlations on all variables. First, we constructed the behavioral model that involved cognitive control as the independent variable, perspective taking as the indirect effect, and total CU traits as the dependent variable. Then we used network density to evaluate hypothesized brain associations as independent variables with associated behavioral data. After that we evaluated between network associations that were suggested by the correlation matrices prior to analysis. We then took all identified models and placed them in one final model and conducted a final set of likelihood ratio tests to arrive at the most parsimonious model.

#### Examining Social and Conflict Networks Connection Density with the dlPFC

We examined associations between CU traits, cognitive control, and perspective taking with connection density between social and conflict networks with the dlPFC using a path analysis controlling for pubertal stage, sex, and age.

#### Node Centrality

To identify central nodes for each network and their associations with CU traits, perspective taking, and cognitive control, we conducted a set of path analyses evaluating centralities for all nodes of the social network in one model and then a model with centrality for all nodes in the conflict network in another model. Full results are in the Supplementary Results.

## 3. Results

### 3.1. Descriptives

Sample and GIMME network descriptives are presented in Supplementary Results and Supplementary Figure 1.

### 3.2. Full Model

Placing individual models during model building (For model building: Supplementary Results and Supplementary Figure 2, Supplemental Tables 1 and 2) into one full model retained most of the parameters. The only parameter we removed was the indirect effect of negative density between social and conflict networks via perspective taking on CU traits (Supplementary Table 2). The final model fit the data (X^2^ 19.193(17), CFI= 0.981, TLI = 0.950; RMSEA= 0.035; SRMR= 0.053; Figure 2 for full model specification, Supplemental Table 3 for model fitting).

**Figure 1.**
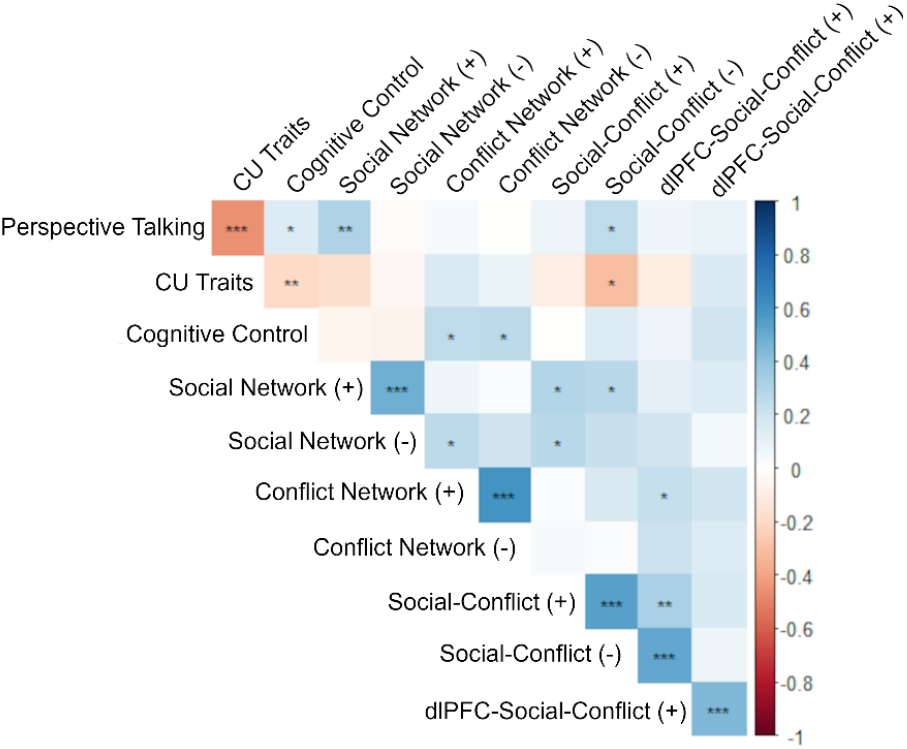
Zero-order correlations of independent variables with brain parameters. Note *= P< 0.05, **= p<0.01, *** p< 0.001, (+) = positive connection density, (-) = negative connection density

**Figure 2.**
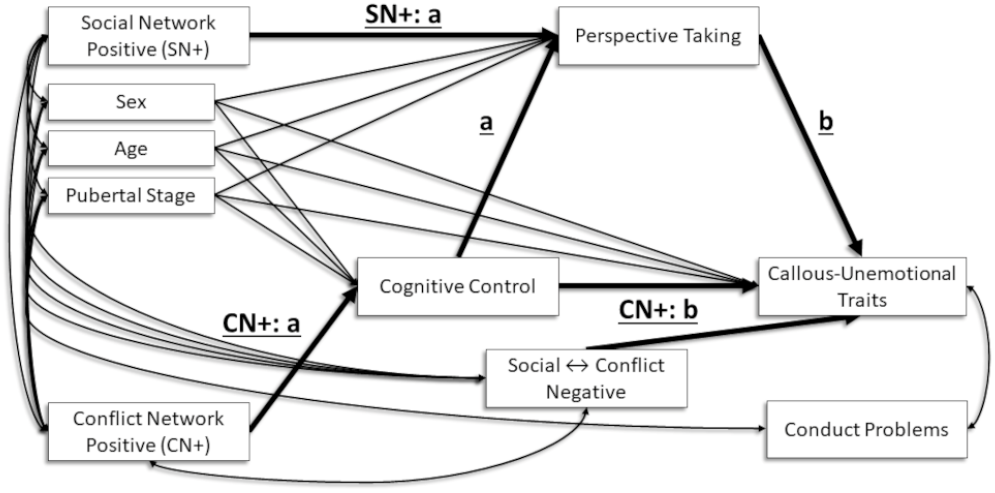
Depicting Full Path Model Specification

**Figure 3.**
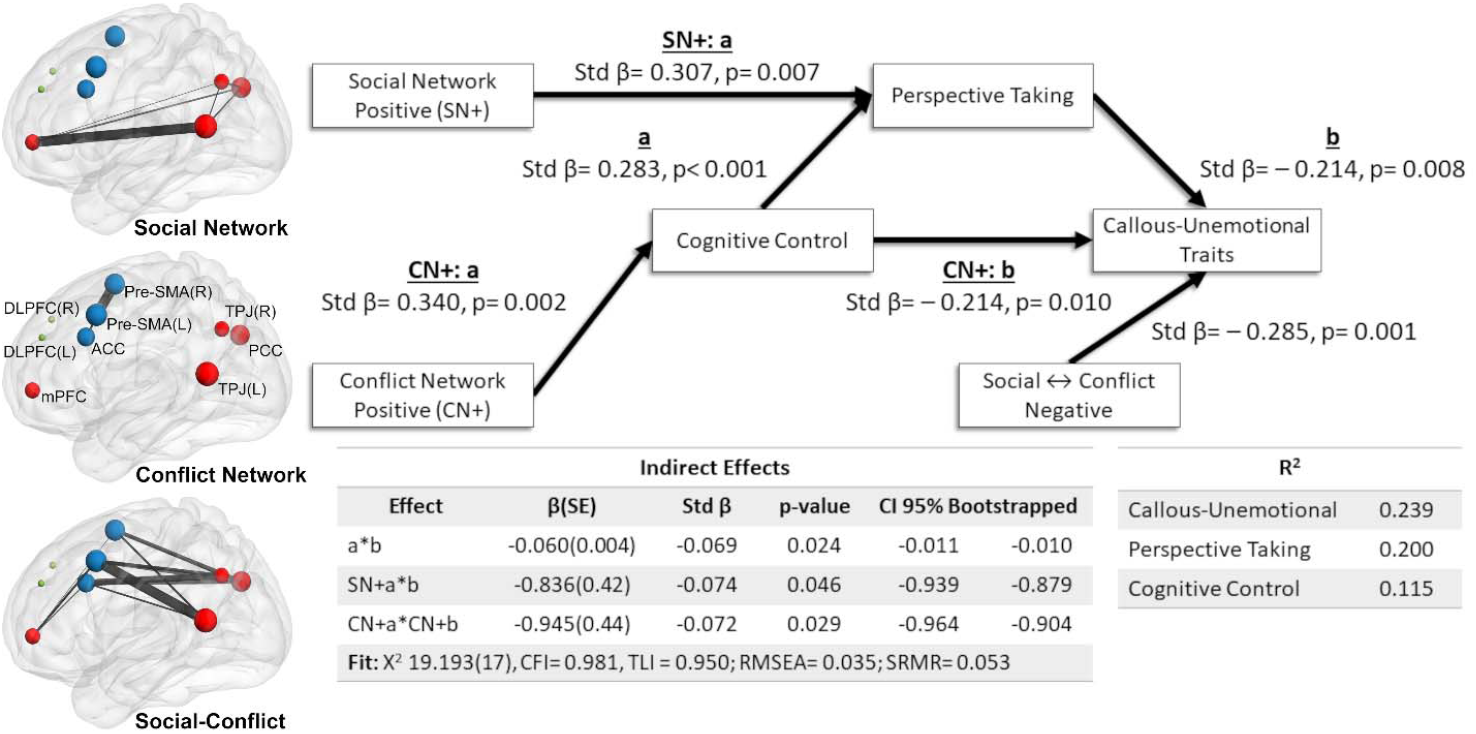
Depicting the Final Full Model Results and Average Functional Network Properties of Centrality and Density

For the behavioral model, Cognitive Control negatively associated with CU traits indirectly via perspective taking (Table 1, Figure 4). For brain associations, positive density in the social network indirectly associated with CU traits via perspective taking with a negative indirect effect; and that positive density in the conflict network indirectly associated with CU traits via Cognitive Control and this indirect effect was negative (Table 1, Figure 4). The negative density between the social and conflict networks only retained a direct effect on CU traits that was negative (Table 1, Figure 4). The model accounted for 24% of the variance in CU traits, 20% of the variance in perspective taking, and 12% of the variance in cognitive control.

**Table 1.** Final Model Results

| | Unstd $\beta$ | SE | Std $\beta$ | z | p-value | 95% CI Bootstrapped | |
| --- | --- | --- | --- | --- | --- | --- | --- |
|  |  |  |  |  |  | Lower | Upper |
| **Callous-Unemotional Traits ~ (R<sup>2</sup>= 0.239)** |  |  |  |  |  |  |  |
| Perspective taking “b” | -0.405* | 0.154 | -0.242 | -2.624 | 0.009 | -0.435 | -0.414 |
| cognitive control “CN+: b” | -0.032* | 0.012 | -0.214 | -2.582 | 0.010 | -0.033 | -0.032 |
| Social ↔ conflict network (-) | -2.572* | 0.809 | -0.285 | -3.177 | 0.001 | -2.628 | -2.514 |
| Tanner | 1.171 | 0.798 | 0.152 | 1.467 | 0.142 | -0.393 | 2.734 |
| Sex | 2.481 | 1.476 | 0.153 | 1.681 | 0.093 | -0.413 | 5.375 |
| Age | 0.115 | 0.551 | 0.019 | 0.209 | 0.835 | -0.965 | 1.195 |
| **Perspective Taking ~ (R<sup>2</sup>= 0.200)** |  |  |  |  |  |  |  |
| cognitive control “a” | 0.025* | 0.006 | 0.283 | 3.941 | 0.000 | 0.024 | 0.025 |
| Social network (+) “SN+: a” | 2.063* | 0.761 | 0.307 | 2.709 | 0.007 | 2.143 | 2.248 |
| Tanner | 0.508 | 0.444 | 0.110 | 1.144 | 0.253 | -0.362 | 1.378 |
| Sex | -1.218 | 0.955 | -0.125 | -1.275 | 0.202 | -3.090 | 0.655 |
| Age | -0.778* | 0.340 | -0.216 | -2.285 | 0.022 | -0.792 | -0.748 |
| **cognitive control ~ (R<sup>2</sup>= 0.115)** |  |  |  |  |  |  |  |
| Conflict network (+) “CN+: a” | 29.959* | 9.625 | 0.340 | 3.113 | 0.002 | 28.901 | 30.168 |
| Tanner | -1.678 | 5.441 | -0.031 | -0.308 | 0.758 | -2.001 | 1.332 |
| Sex | 2.178 | 2.400 | 0.085 | 0.907 | 0.364 | -2.059 | 2.363 |
| Age | -1.678 | 5.441 | -0.031 | -0.308 | 0.758 | -4.755 | 5.515 |
| **Indirect Effects** |  |  |  |  |  |  |  |
| a*b | -0.010* | 0.004 | -0.069 | -2.261 | 0.024 | -0.011 | -0.010 |
| SN+: a*b | -0.836* | 0.419 | -0.074 | -1.993 | 0.046 | -0.939 | -0.879 |
| CN+: a* CN+: b | -0.945* | 0.436 | -0.073 | -2.168 | 0.030 | -0.964 | -0.904 |

Note: bias-corrected bootstrapped confidence intervals with 5000 resamples
\* = $p < 0.05$

### 3.3. Associations of Social and Conflict Networks Communicating with dlPFC

Although zero-order correlations did not detect a relationship (Figure 1), follow up analysis controlling for pubertal stage, sex, and age revealed cognitive control (Std-β= 0.323, p= 0.001) and puberty (Std-β= 0.321, p= 0.018) positively associated with negative density of the dlPFC with social and conflict networks, whereas age negatively associated (Std-β= −0.208, p= 0.041; Table 2).

**Table 2.** Network Associations with the DLPFC

| | Unstd $\beta$ | SE | Std $\beta$ | z | p-value | 95% CI Bootstrapped | |
| --- | --- | --- | --- | --- | --- | --- | --- |
|  |  |  |  |  |  | Lower | Lower |
| **Negative Social and Conflict with DLPFC ~ (R<sup>2</sup>= 0.220)** |  |  |  |  |  |  |  |
| Callous-Unemotional Traits | 0.024 | 0.017 | 0.220 | 1.445 | 0.149 | -0.008 | 0.056 |
| Perspective Taking | 0.026 | 0.021 | 0.141 | 1.230 | 0.219 | -0.015 | 0.068 |
| Cognitive Control | 0.005* | 0.002 | 0.323 | 3.205 | 0.001 | 0.002 | 0.008 |
| Tanner | 0.269* | 0.113 | 0.321 | 2.374 | 0.018 | 0.047 | 0.490 |
| Sex | -0.374 | 0.191 | -0.210 | -1.961 | 0.050 | -0.748 | -0.002 |
| Age | -0.137 | 0.067 | -0.208 | -2.047 | 0.041 | -0.268 | -0.006 |
| **Positive Social and Conflict with DLPFC ~ (R<sup>2</sup>= 0.093)** |  |  |  |  |  |  |  |
| Callous-Unemotional Traits | -0.014 | 0.013 | -0.136 | -1.120 | 0.263 | -0.039 | 0.011 |
| Perspective Taking | 0.004 | 0.022 | 0.020 | 0.169 | 0.866 | -0.039 | 0.045 |
| Cognitive Control | 0.001 | 0.002 | 0.051 | 0.435 | 0.664 | -0.003 | 0.004 |
| Tanner | 0.206 | 0.143 | 0.252 | 1.442 | 0.149 | -0.074 | 0.486 |
| Sex | 0.004 | 0.237 | 0.003 | 0.018 | 0.985 | -0.459 | 0.468 |
| Age | 0.014 | 0.121 | 0.022 | 0.116 | 0.908 | -0.223 | 0.251 |

Note: bias-corrected bootstrapped confidence intervals with 5000 resamples
\* = $p < 0.05$

### 3.4. Central Nodes Identified for Each Network

In the social network, perspective taking associated positively with node centrality in the right TPJ (Std-β= 0.334, p= 0.007). We found no associations with variables of interest for centrality within the conflict network. Between social and conflict networks analyses indicate that perspective taking associated with centrality of the ACC (Std-β= 0.314, p= 0.007). The dlPFC connection with social and conflict networks indicated that perspective taking positively associated with centrality with the right TPJ (Std-β= 0.393, p= 0.001) and negatively with mPFC (Std-β= −0.265, p= 0.018; full results in Supplemental Tables 5 through 8).

## 4. Discussion

Results from a community sample of adolescents demonstrates cognitive control is important for perspective taking in relation to CU traits. Additionally, positive connection density in the social network indirectly associated with CU traits via perspective taking and, similarly, the conflict network indirectly associated with CU traits via cognitive control; and less negative density between these networks directly associated with CU traits. This indicates that (1) greater cohesion in the social network associates with higher perspective tasking and mitigates CU traits whereas (2) greater cohesion in the conflict network associates with higher cognitive control and mitigates CU traits, and finally (3) less functional coupling between the social and conflict networks (i.e., one active while the other is inactive) mitigated CU traits directly. In other words, positive connection density within the social and conflict networks as well as functional coupling between the social and conflict networks (i.e., more negative between-network density) could mitigate CU traits in adolescents. Moreover, brain associations were heterogenous indicating the importance of considering individual-specific alterations in neural circuitry. The present results evidence the importance of cognitive control for perspective taking in CU traits and identifies heterogenous functional properties of adolescent brains important for understanding these processes.

### 4.1. Cognitive Control Impacts Callous-Unemotional Traits Via Perspective Taking

Cognitive control indirectly associated with CU traits via perspective taking and the overall indirect effect was negative. Multiple studies converge on cognitive control as an important component of perspective taking (for reviews see: Mahy et al., 2014; Wade et al., 2018); while the CU trait literature demonstrates impairments in cognitive control (Baskin-Sommers et al., 2015; Gluckman et al., 2016; Zeier et al., 2012), perspective taking (Anastassiou-Hadjicharalambous & Warden, 2008; Lui et al., 2016; O’Kearney et al., 2017), and that cognitive control impacts the capacity to infer complex emotions (Winters & Sakai, 2021). The present results extend this literature by demonstrating an overall negative indirect effect of cognitive control on CU trait symptoms via perspective taking. One plausible interpretation is that the automaticity of perspective taking is compromised and may be driven by impairments in cognitive control in adolescents at higher CU traits. This is particularly plausible given that psychopathic adults have less automaticity for taking others perspective (Drayton et al., 2018) and they share the same neurocognitive impairments in youth higher in CU traits (Viding & McCrory, 2012). The behavioral component of this model connects the extant literature and provides novel evidence that cognitive control influences perspective taking in relation to CU traits.

### 4.2. Social Network Impacts Callous-Unemotional Traits Via Perspective Taking

Positive density in the social network had a negative indirect effect on CU traits via perspective taking, which may indicate that functional integration of the social network, and the extent to which this occurs, is important for CU traits. This effect may be gated by the TPJ as greater node centrality in the right TPJ was positively associated with perspective taking. The social network is known to support perspective taking (Blakemore, 2012; Klapwijk et al., 2013; McCormick et al., 2018) and regions comprising this social network are less functionally integrated in the presence of CU traits (Thijssen & Kiehl, 2017; Umbach & Tottenham, 2020). While previous research speculated that lower functional integration of these regions may underlie social impairments observed in CU traits, we demonstrate that positive network connection density in the social network is important for perspective taking and reduced CU traits. Although we cannot make causal interpretations with the present data, the negative indirect effect in the present study along with the extant literature suggests that improving functional integration of the social network may increase perspective taking capacity (Kral et al., 2017; Winters, Pruitt, et al., 2021), which may mitigate CU trait symptoms. Behaviorally this may look like a greater capacity to monitor the environment and resolving conflicts between self and others’ emotions that leads to improved emotional responsiveness. Thus, this finding warrants further investigation directly testing the social networks involvement in perspective taking in CU trait symptoms, particularly in the right TPJ.

### 4.3. Conflict Network Impacts Callous-Unemotional Traits Via Cognitive Control

Positive connection density in the conflict network positively associated with cognitive control, which was consistent with previous task-based literature (e.g., Iannaccone, Hauser, Ball, et al., 2015; Iannaccone, Hauser, Staempfli, et al., 2015). Surprisingly, the conflict network had no direct association with CU traits, but rather indirectly associated with CU traits via cognitive control. While previous literature demonstrates aberrant function of these regions in CU traits (e.g., Pu et al., 2017; Szabó et al., 2017; Yoder et al., 2016), the present finding suggests this may be accounted for by cognitive control. This is plausible given that this network is strongly associated with tasks involving conflict (e.g., Banich, 2019; Iannaccone, Hauser, Ball, et al., 2015; Iannaccone, Hauser, Staempfli, et al., 2015; Milham & Banich, 2005). The negative indirect effect suggests greater functional integration of the conflict network could improve cognitive control and mitigate CU traits. Importantly, the conflict networks association with cognitive control that associates with perspective taking and indirectly mitigates CU traits is a novel path indicating the brain’s involvement in cognitive control facilitates the cognitive control necessary for responding to others’ emotions. This supports findings that conflict network nodes are critical for resolving conflict regarding others emotions (Maier & di Pellegrino, 2012). Thus, functional integration within this network could indirectly mitigate CU traits.

### 4.4. Conflict Network Node Centrality Not Associated with Cognitive Control

Our hypothesis that cognitive control would associate with node centrality in the ACC of the conflict network was not supported. This suggests the importance of distributed function, as opposed to node centrality, in the conflict network amongst adolescents.

### 4.5. Between Social and Conflict Network Density and Callous-Unemotional Traits

Greater negative density between the social and conflict networks directly associated with lower CU traits. This is consistent with literature on adult psychopathy (Dotterer et al., 2020) and adolescent CU traits (Pu et al., 2017; Winters, Sakai, et al., 2021) demonstrating that weaker anticorrelation between task positive and task negative networks associate with CU traits. Given that anticorrelation between task positive and negative networks indicates greater maturity (Richardson et al., 2018), it is plausible that there is a developmental delay in underlying neural circuitry associated with CU traits; which would support the notion that psychopathic traits involve a delay in neurodevelopment (Frick & Viding, 2009). Overall, this finding adds to the mounting evidence that there are differences in functional brain coupling between task positive and negative networks, which may be indicative of developmental immaturity, amongst those higher in CU traits.

### 4.6. Network Density with dlPFC Associates with Cognitive Control

Present findings partially support previous literature indicating the dlPFC, along with the conflict network, exerts top-down control over the social network when resolving conflict during perspective taking. Cognitive control positively associated with negative density between these networks suggesting some top-down control of the social network but, contrary to expectations, there was no association with perspective taking. However, perspective taking significantly accounted for central nodes between these networks – specifically perspective taking associated positively with the right TPJ and negatively with the mPFC. This suggests that although density with the dlPFC was not associated, greater centrality in the right TPJ and less centrality in the mPFC characterize the dlPFC relationship with the social network related to perspective taking. Together, this suggests that centrality of the right TPJ in communication with the dlPFC is important for perspective taking and that density with the dlPFC to these nodes are important for cognitive control. This could presumably indicate information processing streams related to top-down control, and suggests that investigating the density of connections to the right TPJ may be of particular importance for understanding conflict resolution during perspective taking.

These results must be considered under some limitations. First, this study was observational and did not employ task manipulations to capture cognitive control’s modulation of perspective taking. Future studies could build on the present finding by using a paradigm that includes different levels of demand on cognitive control during perspective taking. Second, this study was cross sectional and cannot make any causal interpretation. The literature provides substantial support for the ordering of variables and interpretations made; however, these results require further investigation with temporal ordering to confirm. Third, the sample size is modest (n= 87) and may have missed some effects. Fourth, we were not able to obtain the granularity required to examine other divisions in the ACC (e.g., de la Vega et al., 2016) that could be addressed in future studies. Fifth, the present sample was a community sample and may not be generalizable to clinical/forensic samples. Finally, cognitive control involves multiple cognitive processes that make it unclear exactly what cognitive process was driving the associations observed. Future directions of this work are to isolate aspects of cognitive control for specific deficits at higher CU traits and determine if granular components or general cognitive control could better explain this relationship.

#### Conclusion

The present study demonstrates the importance of cognitive control for perspective taking in relation to CU traits and identifies the associated functional brain properties relevant for future investigation. The S-GIMME approach improves connection estimation beyond traditional methods by accounting for contemporaneous, lagged, and directional connections of individual connectivity maps, which *b*olsters statistical inferences with functional brain properties. Functional brain properties of the social and conflict networks demonstrated indirect associations with CU traits via perspective taking and cognitive control (respectively) that are important considerations for cognitive controls influence on perspective taking for CU traits. These novel findings *support* the importance of the relationship between cognitive control and socio-cognitive processes in CU traits, *while extending* this work by demonstrating the importance of integrated brain regions supporting these processes. Together this model characterizes a complicated set of processes and underlying functional brain properties underlying CU traits in adolescents. Future studies could build on current findings by modeling the heterogenous brain features of networks specific to these processes found here and further probing these associations in behavioral paradigms with task manipulations to isolate aspects of cognitive control. Taking account of the individual heterogeneity of adolescent brains could be leveraged to improve individualized treatment approaches for these youth.

