## Supplemental for "Resting-state connectivity underlying cognitive control’s association with perspective taking in callous-unemotional traits"

**Supplementary Material**

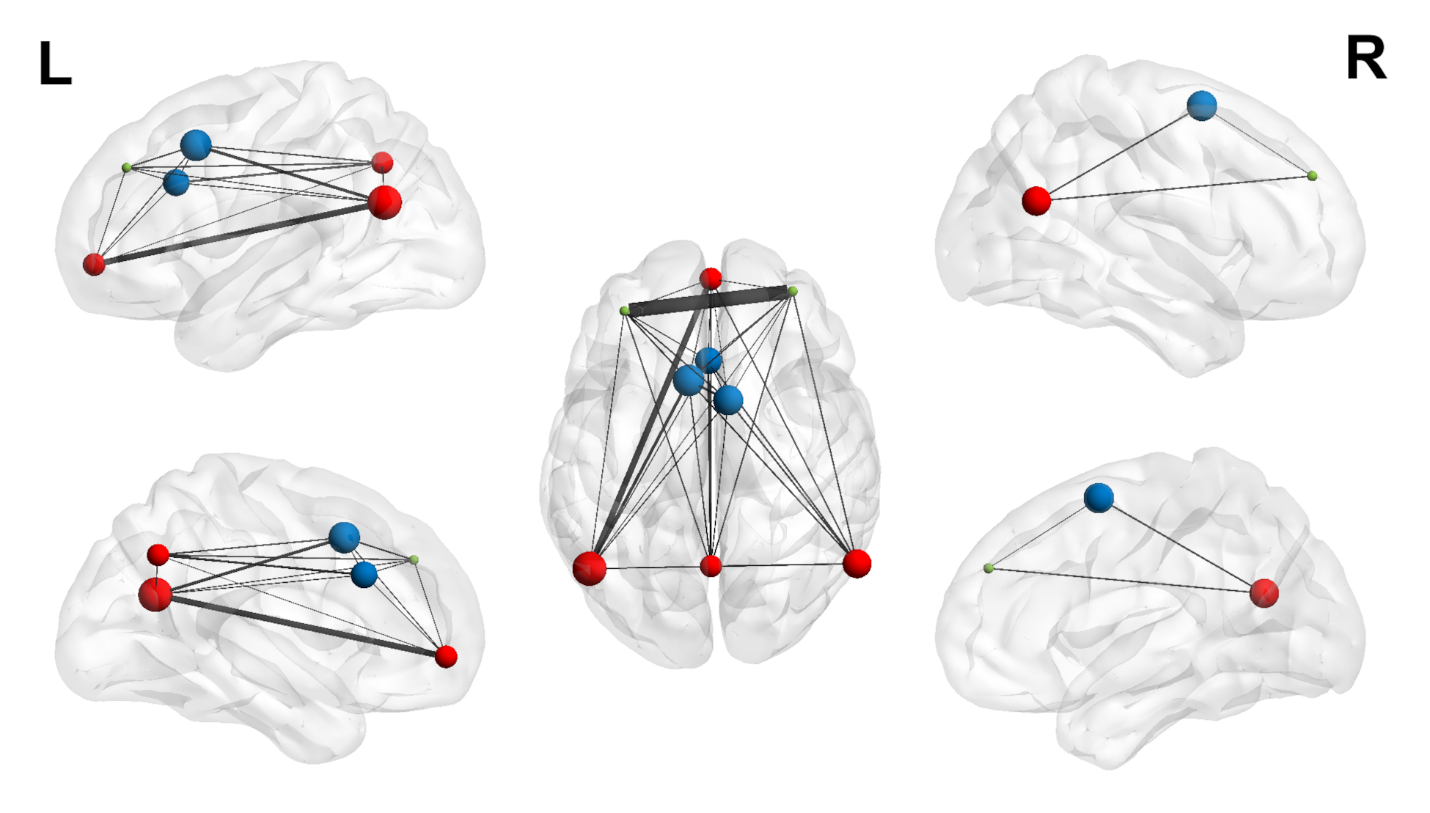

Supplementary Figure 1. Average brain centrality and connection density across all participants

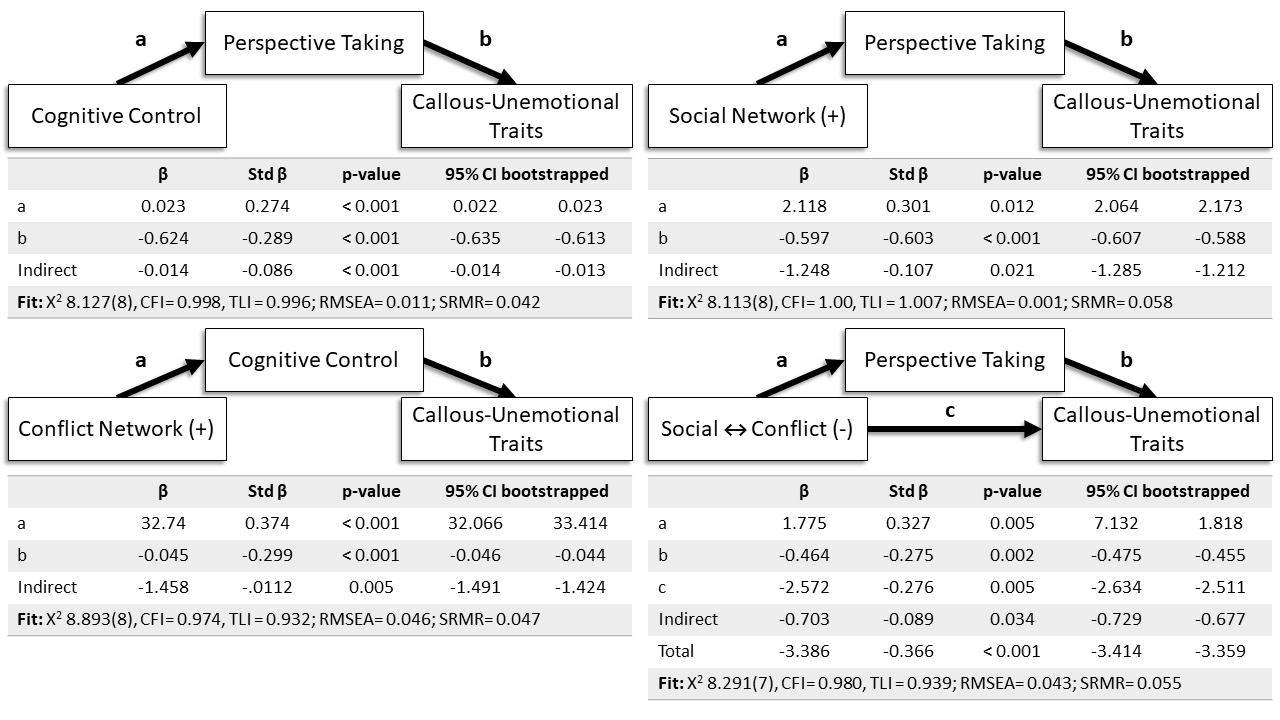

Supplementary Figure 1. Individual Model Building Results

| Supplemental Table 1. Individual Model Building Likelihood Ratio Tests | | | | | | | | |
| --- | --- | --- | --- | --- | --- | --- | --- | --- |
|  | BIC | X^2^(df) | CFI | TLI | RMSEA | ΔX^2^ (df) | ΔX^2^ p | Decision |
| **Behavior model** |  |  |  |  |  |  |  |  |
| Individual Model |  |  |  |  |  |  |  |  |
| Unconstrained model | 4297 | 4.739 (7) | 1.00 | 1.107 | 0.000 |  |  |  |
| Constraining indirect | 4299 | 8.601(8) | 0.942 | 0.847 | 0.066 | 12.048(1) | < 0.001 | Retain indirect |
| Constraining direct | 3296 | 8.126 (8) | 0.998 | 0.996 | 0.011 | 3.780(1) | 0.052 | Remove direct |
| **Social network (+)** |  |  |  |  |  |  |  |  |
| Individual Model |  |  |  |  |  |  |  |  |
| Unconstrained model | 3067 | 7.709(7) | 0.993 | 0.978 | 0.024 |  |  |  |
| Constraining indirect | 3038 | 13.164(8) | 0.919 | 0.787 | 0.076 | 5.411(1) | 0.020 | Retain indirect |
| Constraining direct | 3063 | 8.112(8) | 1.000 | 1.007 | 0.001 | 0.403(1) | 0.525 | Remove direct |
| **Conflict Network (+)** |  |  |  |  |  |  |  |  |
| Individual Model |  |  |  |  |  |  |  |  |
| Unconstrained model | 3540 | 6.754(7) | 0.992 | 0.975 | 0.028 |  |  |  |
| Constraining indirect | 3539 | 10.339(8) | 0.945 | 0.857 | 0.067 | 8.732(1) | 0.003 | Retain indirect |
| Constraining direct | 3539 | 8.893((8) | 0.974 | 0.932 | 0.046 | 2.319(1) | 0.611 | Remove direct |
| **Between social ↔ conflict networks (-)** | | |  |  |  |  |  |  |
| Individual Model |  |  |  |  |  |  |  |  |
| Unconstrained model | 3090 | 8.291(7) | 0.980 | 0.939 | 0.043 |  |  |  |
| Constraining indirect | 3093 | 15.231(8) | 0.892 | 0.715 | 0.093 | 7.226(1) | 0.007 | Retain indirect |
| Constraining direct | 3091 | 13.073(8) | 0.923 | 0.797 | 0.079 | 5.197(1) | 0.023 | Retain direct |
| *= given the marginal significance – we investigated the beta coefficients to determine if it significantly contributed to the model. Because the beta coefficient and the likelihood ratio test were insignificant – we removed this indirect effect for the final model. | | | | | | | | |

| Supplemental Table 2. Final Model Building Likelihood Ratio Tests | | | | | | | | |
| --- | --- | --- | --- | --- | --- | --- | --- | --- |
|  | BIC | X^2^(df) | CFI | TLI | RMSEA | ΔX^2^ (df) | ΔX^2^ p | Decision |
| Unconstrained model | 4297 | 4.739 (7) | 1.00 | 1.107 | 0.000 |  |  |  |
| **Behavior model** |  |  |  |  |  |  |  |  |
| Constraining indirect | 4545 | 24.360(16) | 0.920 | 0.774 | 0.074 | 6.555 (1) | 0.010 | Retain indirect |
| **Social network (+)** |  |  |  |  |  |  |  |  |
| Constraining indirect | 4544 | 22.812(16) | 0.933 | 0.812 | 0.067 | 4.785 (1) | 0.028 | Retain indirect |
| **Conflict Network (+)** |  |  |  |  |  |  |  |  |
| Constraining indirect | 4546 | 25.063(16) | 0.910 | 0.746 | 0.780 | 15.232(1) | < 0.001 | Retain indirect |
| **Between social ↔ conflict networks (-)** | | |  |  |  |  |  |  |
| Constraining indirect | 4543 | 21.973(16) | 0.953 | 0.869 | 0.051 | 3.985 (1) | 0.050 | Remove indirect* |
| Constraining direct | 4544 | 22.848(16) | 0.933 | 0.812 | 0.0.67 | 4.620(1) | 0.032 | Retain direct |
| *= given the marginal significance – we investigated the beta coefficients to determine if it significantly contributed to the model. Because the beta coefficient and the likelihood ratio test were insignificant – we removed this indirect effect for the final model. | | | | | | | | |

| Supplemental Table 3. Fit for final models ran initially and after correlating error terms | | | | | |
| --- | --- | --- | --- | --- | --- |
|  | X^2^(df) | CFI | TLI | RMSEA | SRMR |
| Base final model | 22.643(17) | 0.947 | 0.861 | 0.056 | 0.058 |
| Correlating puberty and conduct residuals | 19.092 (16) | 0.973 | 0.925 | 0.043 | 0.053 |

| Supplemental Table 4. Multigroup model | | | | | |
| --- | --- | --- | --- | --- | --- |
| Model | BIC | X^2^(df) | ΔX^2^ (df) | ΔX^2^ p | Decision |
| Unconstrained – grouped by sex | 4471 | 34.218(35) |  |  |  |
| Constrained – regression parameters by sex | 4454 | 35.532(39) | 1.531(4) | 0.821 | No Difference |

| Supplemental Table 5. Centrality within the Social Network | | | | | |
| --- | --- | --- | --- | --- | --- |
|  | Unstd β | SE | Std β | z | p-value |
| **TPJ(R) ~ (R^2^= 0.115)** | | |  |  |  |
| Callous-Unemotional Traits | 0.003 | 0.002 | 0.171 | 1.219 | 0.223 |
| Perspective Taking | 0.010* | 0.003 | 0.334 | 2.698 | 0.007 |
| Conflict Adaptation | 0.000 | 0.000 | -0.127 | -1.032 | 0.302 |
| Tanner | -0.003 | 0.018 | -0.02 | -0.144 | 0.885 |
| Sex | 0.038 | 0.031 | 0.142 | 1.221 | 0.222 |
| Age | 0.010 | 0.013 | 0.105 | 0.803 | 0.422 |
| **TPJ(L) ~ (R^2^= 0.110)** | | |  |  |  |
| Callous-Unemotional Traits | -0.001 | 0.002 | -0.088 | -0.641 | 0.521 |
| Perspective Taking | -0.007 | 0.004 | -0.255 | -1.835 | 0.067 |
| Conflict Adaptation | 0.000 | 0.000 | -0.009 | -0.062 | 0.951 |
| Tanner | 0.024 | 0.020 | 0.184 | 1.211 | 0.226 |
| Sex | -0.073* | 0.033 | -0.261 | -2.209 | 0.027 |
| Age | -0.006 | 0.014 | -0.054 | -0.408 | 0.683 |
| **mPFC ~ (R^2^= 0.115)** | | |  |  |  |
| Callous-Unemotional Traits | -0.001 | 0.001 | -0.076 | -0.570 | 0.569 |
| Perspective Taking | 0.000 | 0.002 | -0.011 | -0.094 | 0.925 |
| Conflict Adaptation | 0.000 | 0.000 | 0.028 | 0.238 | 0.812 |
| Tanner | -0.003 | 0.010 | -0.034 | -0.270 | 0.787 |
| Sex | -0.039 | 0.021 | -0.221 | -1.905 | 0.057 |
| Age | 0.016 | 0.009 | 0.24 | 1.724 | 0.085 |
| **PCC ~ (R^2^= 0.167)** | | |  |  |  |
| Callous-Unemotional Traits | -0.003 | 0.001 | -0.233 | -1.736 | 0.083 |
| Perspective Taking | 0.003 | 0.002 | 0.173 | 1.575 | 0.115 |
| Conflict Adaptation | 0.000 | 0.000 | -0.164 | -1.487 | 0.137 |
| Tanner | 0.018 | 0.010 | 0.217 | 1.723 | 0.085 |
| Sex | -0.006 | 0.020 | -0.035 | -0.302 | 0.762 |
| Age | 0.006 | 0.007 | 0.089 | 0.822 | 0.411 |
| Note: *= p < 0.05 | | | | | |

| Supplemental Table 6. Centrality within the Conflict Network | | | | | |
| --- | --- | --- | --- | --- | --- |
|  | Unstd β | SE | Std β | z | p-value |
| **ACC ~ (R^2^= 0.230)** | | |  |  |  |
| Callous-Unemotional Traits | 0.002 | 0.001 | 0.240 | 1.802 | 0.072 |
| Perspective Taking | 0.004 | 0.002 | 0.273 | 1.675 | 0.094 |
| Conflict Adaptation | 0.000 | 0.000 | 0.063 | 0.656 | 0.512 |
| Tanner | 0.001 | 0.007 | 0.018 | 0.157 | 0.875 |
| Sex | -0.018 | 0.015 | -0.134 | -1.218 | 0.223 |
| Age | 0.017* | 0.006 | 0.343 | 2.875 | 0.004 |
| **Pre-SMA(R) ~ (R^2^= 0.152)** | | |  |  |  |
| Callous-Unemotional Traits | 0.004 | 0.002 | 0.242 | 1.735 | 0.083 |
| Perspective Taking | 0.002 | 0.003 | 0.090 | 0.746 | 0.456 |
| Conflict Adaptation | 0.000 | 0.000 | 0.198 | 1.592 | 0.111 |
| Tanner | -0.007 | 0.021 | -0.057 | -0.327 | 0.743 |
| Sex | 0.022 | 0.030 | 0.087 | 0.732 | 0.464 |
| Age | 0.023 | 0.015 | 0.243 | 1.488 | 0.137 |
| **Pre-SMA(L) ~ (R^2^= 0.173)** | | |  |  |  |
| Callous-Unemotional Traits | 0.001 | 0.002 | 0.056 | 0.360 | 0.719 |
| Perspective Taking | -0.001 | 0.003 | -0.021 | -0.157 | 0.876 |
| Conflict Adaptation | 0.001* | 0.000 | 0.242 | 2.017 | 0.044 |
| Tanner | -0.021 | 0.019 | -0.184 | -1.072 | 0.284 |
| Sex | -0.046 | 0.029 | -0.192 | -1.589 | 0.112 |
| Age | 0.024 | 0.013 | 0.270 | 1.809 | 0.070 |
| Note: *= p < 0.05 | | | | | |

| Supplemental Table 7. Centrality Between the Social and Conflict Networks | | | | | |
| --- | --- | --- | --- | --- | --- |
|  | Unstd β | SE | Std β | z | p-value |
| **ACC ~ (R^2^= 0.115)** | | |  |  |  |
| Callous-Unemotional Traits | 0.001 | 0.002 | 0.065 | 0.476 | 0.634 |
| Perspective Taking | 0.006* | 0.002 | 0.314 | 2.723 | 0.006 |
| Conflict Adaptation | 0.000 | 0.000 | -0.108 | -0.846 | 0.398 |
| Tanner | -0.001 | 0.012 | -0.016 | -0.121 | 0.904 |
| Sex | 0.042 | 0.022 | 0.228 | 1.907 | 0.057 |
| Age | 0.006 | 0.009 | 0.087 | 0.656 | 0.512 |
| **SMA(R) ~ (R^2^= 0.071)** | | |  |  |  |
| Callous-Unemotional Traits | 0.000 | 0.001 | -0.051 | -0.342 | 0.732 |
| Perspective Taking | 0.000 | 0.002 | -0.018 | -0.134 | 0.893 |
| Conflict Adaptation | 0.000 | 0.000 | -0.083 | -0.783 | 0.434 |
| Tanner | 0.006 | 0.011 | 0.089 | 0.586 | 0.558 |
| Sex | -0.005 | 0.017 | -0.032 | -0.271 | 0.786 |
| Age | 0.012 | 0.008 | 0.216 | 1.472 | 0.141 |
| **SMA(L) ~ (R^2^= 0.165)** | | |  |  |  |
| Callous-Unemotional Traits | -0.001 | 0.001 | -0.074 | -0.472 | 0.637 |
| Perspective Taking | 0.001 | 0.001 | 0.060 | 0.544 | 0.586 |
| Conflict Adaptation | 0.000 | 0.000 | -0.036 | -0.319 | 0.750 |
| Tanner | 0.002 | 0.009 | 0.031 | 0.219 | 0.826 |
| Sex | -0.008 | 0.014 | -0.064 | -0.583 | 0.560 |
| Age | 0.018* | 0.006 | 0.380 | 3.322 | 0.001 |
| **TPJ(R) ~ (R^2^= 0.138)** | | |  |  |  |
| Callous-Unemotional Traits | -0.002 | 0.001 | -0.226 | -1.589 | 0.112 |
| Perspective Taking | 0.001 | 0.002 | 0.080 | 0.584 | 0.559 |
| Conflict Adaptation | 0.000 | 0.000 | 0.025 | 0.254 | 0.799 |
| Tanner | 0.012 | 0.009 | 0.196 | 1.326 | 0.185 |
| Sex | -0.018 | 0.017 | -0.137 | -1.037 | 0.300 |
| Age | 0.002 | 0.008 | 0.047 | 0.291 | 0.771 |
| **TPJ(L) ~ (R^2^= 0.122)** | | |  |  |  |
| Callous-Unemotional Traits | 0.000 | 0.001 | -0.037 | -0.261 | 0.794 |
| Perspective Taking | 0.003 | 0.002 | 0.204 | 1.560 | 0.119 |
| Conflict Adaptation | 0.000 | 0.000 | -0.044 | -0.443 | 0.658 |
| Tanner | 0.000 | 0.010 | 0.007 | 0.042 | 0.967 |
| Sex | -0.001 | 0.014 | -0.008 | -0.073 | 0.942 |
| Age | 0.013* | 0.006 | 0.292 | 2.096 | 0.036 |
| **mPFC ~ (R^2^= 0.065)** | | |  |  |  |
| Callous-Unemotional Traits | 0.000 | 0.001 | -0.006 | -0.043 | 0.966 |
| Perspective Taking | 0.001 | 0.002 | 0.052 | 0.385 | 0.700 |
| Conflict Adaptation | 0.000 | 0.000 | -0.018 | -0.114 | 0.909 |
| Tanner | 0.000 | 0.009 | 0.005 | 0.032 | 0.975 |
| Sex | 0.007 | 0.016 | 0.058 | 0.463 | 0.644 |
| Age | 0.012 | 0.007 | 0.249 | 1.587 | 0.113 |
| **PCC ~ (R^2^= 0.054)** | | |  |  |  |
| Callous-Unemotional Traits | 0.001 | 0.001 | 0.088 | 0.622 | 0.534 |
| Perspective Taking | -0.002 | 0.002 | -0.121 | -0.952 | 0.341 |
| Conflict Adaptation | 0.000 | 0.000 | 0.107 | 0.797 | 0.425 |
| Tanner | -0.004 | 0.007 | -0.061 | -0.512 | 0.609 |
| Sex | -0.011 | 0.017 | -0.088 | -0.679 | 0.497 |
| Age | 0.006 | 0.007 | 0.136 | 0.988 | 0.323 |
| Note: *= p < 0.05 | | | | | |

| Supplemental Table 8. Centrality Between dlPFC and the Social and Conflict Networks | | | | | |
| --- | --- | --- | --- | --- | --- |
|  | Unstd β | SE | Std β | z | p-value |
| **ACC ~ (R^2^= 0.143)** | | |  |  |  |
| Callous-Unemotional Traits | 0.000 | 0.001 | 0.035 | 0.240 | 0.810 |
| Perspective Taking | 0.002 | 0.002 | 0.159 | 1.134 | 0.257 |
| Conflict Adaptation | 0.000 | 0.000 | -0.077 | -0.671 | 0.502 |
| Tanner | 0.007 | 0.008 | 0.122 | 0.844 | 0.399 |
| Sex | -0.007 | 0.012 | -0.065 | -0.589 | 0.556 |
| Age | 0.011* | 0.005 | 0.275 | 2.217 | 0.027 |
| **SMA(R) ~ (R^2^= 0.049)** | | |  |  |  |
| Callous-Unemotional Traits | 0.000 | 0.001 | -0.009 | -0.079 | 0.937 |
| Perspective Taking | 0.000 | 0.001 | -0.051 | -0.422 | 0.673 |
| Conflict Adaptation | 0.000 | 0.000 | 0.136 | 1.239 | 0.215 |
| Tanner | -0.003 | 0.005 | -0.085 | -0.658 | 0.510 |
| Sex | -0.012 | 0.010 | -0.154 | -1.199 | 0.230 |
| Age | 0.002 | 0.004 | 0.052 | 0.415 | 0.678 |
| **SMA(L) ~ (R^2^= 0.048)** | | |  |  |  |
| Callous-Unemotional Traits | 0.000 | 0.001 | -0.016 | -0.142 | 0.887 |
| Perspective Taking | 0.001 | 0.001 | 0.062 | 0.515 | 0.607 |
| Conflict Adaptation | 0.000 | 0.000 | 0.098 | 0.689 | 0.491 |
| Tanner | 0.002 | 0.009 | 0.027 | 0.164 | 0.870 |
| Sex | -0.001 | 0.015 | -0.007 | -0.053 | 0.958 |
| Age | 0.006 | 0.007 | 0.144 | 0.918 | 0.359 |
| **TPJ(R) ~ (R^2^= 0.074)** | | |  |  |  |
| Callous-Unemotional Traits | 0.001 | 0.001 | 0.098 | 0.588 | 0.557 |
| Perspective Taking | 0.000 | 0.002 | -0.016 | -0.104 | 0.918 |
| Conflict Adaptation | 0.000 | 0.000 | 0.042 | 0.390 | 0.697 |
| Tanner | 0.015 | 0.009 | 0.232 | 1.746 | 0.081 |
| Sex | -0.016 | 0.014 | -0.114 | -1.081 | 0.280 |
| Age | 0.001 | 0.006 | 0.021 | 0.171 | 0.864 |
| **TPJ(L) ~ (R^2^= 0.181)** | | |  |  |  |
| Callous-Unemotional Traits | 0.001 | 0.001 | 0.104 | 0.829 | 0.407 |
| Perspective Taking | 0.005* | 0.002 | 0.393 | 3.297 | 0.001 |
| Conflict Adaptation | 0.000 | 0.000 | -0.04 | -0.321 | 0.749 |
| Tanner | -0.008 | 0.009 | -0.122 | -0.915 | 0.360 |
| Sex | 0.037* | 0.015 | 0.27 | 2.442 | 0.015 |
| Age | 0.010 | 0.006 | 0.189 | 1.655 | 0.098 |
| **mPFC ~ (R^2^= 0.217)** | | |  |  |  |
| Callous-Unemotional Traits | 0.000 | 0.001 | 0.005 | 0.039 | 0.969 |
| Perspective Taking | -0.003* | 0.001 | -0.265 | -2.361 | 0.018 |
| Conflict Adaptation | 0.000 | 0.000 | 0.138 | 1.330 | 0.183 |
| Tanner | 0.005 | 0.007 | 0.116 | 0.822 | 0.411 |
| Sex | -0.019 | 0.010 | -0.19 | -1.848 | 0.065 |
| Age | 0.010 | 0.004 | 0.268 | 2.281 | 0.023 |
| **PCC ~ (R^2^= 0.163)** | | |  |  |  |
| Callous-Unemotional Traits | -0.001 | 0.001 | -0.118 | -0.808 | 0.419 |
| Perspective Taking | 0.002 | 0.001 | 0.151 | 1.130 | 0.259 |
| Conflict Adaptation | 0.000 | 0.000 | 0.058 | 0.676 | 0.499 |
| Tanner | 0.015* | 0.006 | 0.313 | 2.389 | 0.017 |
| Sex | -0.013 | 0.012 | -0.129 | -1.081 | 0.280 |
| Age | 0.000 | 0.005 | -0.006 | -0.046 | 0.963 |
| **mPFC ~ (R^2^= 0.132)** | | |  |  |  |
| Callous-Unemotional Traits | 0.000 | 0.001 | -0.022 | -0.196 | 0.845 |
| Perspective Taking | 0.000 | 0.001 | 0.042 | 0.340 | 0.734 |
| Conflict Adaptation | 0.000 | 0.000 | -0.205 | -1.487 | 0.137 |
| Tanner | 0.006 | 0.008 | 0.166 | 0.779 | 0.436 |
| Sex | 0.007 | 0.009 | 0.083 | 0.705 | 0.481 |
| Age | 0.007 | 0.004 | 0.219 | 1.468 | 0.142 |
| **PCC ~ (R^2^= 0.096)** | | |  |  |  |
| Callous-Unemotional Traits | 0.000 | 0.001 | -0.022 | -0.172 | 0.863 |
| Perspective Taking | 0.000 | 0.001 | -0.031 | -0.210 | 0.833 |
| Conflict Adaptation | 0.000 | 0.000 | 0.185 | 1.745 | 0.081 |
| Tanner | 0.005 | 0.006 | 0.111 | 0.765 | 0.444 |
| Sex | 0.011 | 0.011 | 0.126 | 1.031 | 0.302 |
| Age | 0.003 | 0.005 | 0.076 | 0.537 | 0.591 |
| Note: *= p < 0.05 | | | | | |

**Supplementary Methods**

**Covariates**. All analyses controlled for sex, age, and pubertal stage. Pubertal stage and sex were measured by the genital and breast development subscales of the Tanner assessment (Petersen et al., 1988; current sample α = .77). Because the variation in timing of puberty when measured by age (about five years , Parent et al., 2003) and hormonal changes during puberty impact behavior via direct effect on the adolescent brain (Cameron, 2004; Dahl, 2004; Sisk & Foster, 2004), we controlled for pubertal stage. Similarly, we controlled for age because, as mentioned above, puberty and age measure related yet distinct constructs and, importantly, cognitive control has demonstrated age-related changes with cognitive control improving with age. It is also important to note that the extent to which cognitive control improves is associated with overall cognitive function (Yagi et al., 2020). Sex was included as a covariate to account for variation in brain associations with CU traits. Prior work supports that males with higher levels of CU traits have more volume, and females less volume, in regions related to affect response (Raschle et al., 2018).

Additionally, to retain the signal of CU traits as the independent variable, we evaluated whether modeling the association between conduct problems and CU traits impacted path estimates. Given that there were no changes because of modeling this association and no concerns for suppression effects (e.g., Hyde et al., 2016; Lozier et al., 2014) we only report on models that include conduct problems correlating with CU traits. Conduct problems often co-occur with CU traits but are distinct and account for different outcomes (e.g., Baskin-Sommers et al., 2015; Herpers et al., 2012; Hyde et al., 2015); thus, in order to address this potential confound, we modeled the association between CU traits and conduct problems. We did this using the raw scores for the externalizing subscale of the Youth Self-Report (Achenbach & Rescorla, 2001) as a covariate, which demonstrates acceptable validity and reliability (Achenbach & Rescorla, 2001), and internally consistent in the present sample (α=.87).

**Imaging acquisition**. Instructions for participants during resting state scanning were to keep their eyes closed without falling asleep. Images were collected with a Siemens TimTrio 3T scanner using a blood oxygen level dependent (BOLD) contrast with an interleaved multiband echo planar imaging (EPI) sequence, which included a functional resting state scan (260 EPI volumes; repetition time (TR) 1400ms; echo time (TE) 30ms; flip angle 65^o^; 64 slices, Field of view (FOV) = 224mm, voxel size 2mm isotropic, duration = 10 minutes) and a magnetization prepared rapid gradient echo (MPRAGE) anatomical image (TR= 1900ms, flip angle 9^o^, 176 slices, FOV= 250mm, voxel size= 1mm isotropic). No scans were removed for T1 stabilization because the Siemens sequence collections images after saturation is received.

**Imaging Preprocessing**. With the raw data, we used the CONN toolbox standard preprocessing pipeline (version 18b; Whitfield-Gabrieli & Nieto-Castanon, 2012) that uses Statistical Parametric Mapping (SPM version 12; Penny et al., 2011). The Artifact Detection Tools (ART; <http://www.nitrc.org/projects/artifact_detect>) flagged motion outliers for correction if framewise displacement > 0.5mm and used spike regression to control for motion outliers. Because a fast multiband sequence was used to collect images, slice timing correction was not applied (Glasser et al., 2013; Wu et al., 2011). Physiologic CSF and white matter noise was regressed out of the BOLD signal using anatomic component-based noise correction method (aCompCor; Whitfield-Gabrieli & Nieto-Castanon, 2012). MPRAGE and EPI images were co-registered and normalized to an MNI template. Images were smoothed using a 6mm Gaussian kernel. Finally, to retain resting state signals, a 0.008 and 0.09Hz bandpass filter was used (Satterthwaite et al., 2013).

During preprocessing we found that 24 participants had motion > 3mm and four had >20% of invalid scans. Because this impacts the integrity of the imaging data, we did not retain the time series of these participants. This left a total of 84 participants with full imaging data and 28 participants (25%) without imaging data.

**S-GIMME**. Network maps were derived for each participant from their individual timeseries in R (Version 4.04; R Core Team, 2021) using the ‘lavaan’ (Rosseel, 2012) and ‘‘GIMME’ (Lane et al., 2021) packages. This is a data-driven sparse modeling approach that iteratively adds network connections and uses LaGrange multipliers (Sörbom, 1989) to assess model fit to retain only the statistically meaningful connections (defined as connections that improve fit for 75% of the sample). The connections retained by this sparse modeling approach minimizes spurious contemporaneous connections generated by saturated models (Gates et al., 2010). Both contemporaneous and lagged connections are modeled simultaneously for the entire sample, subgroups (data-derived groups based on functional connections), and individuals (individual-specific connections), which results in a unified structural equation model for each participant (uSEM; Gates et al., 2011). Each participants’ network of connections is derived by iteratively adding connections, assessing fit of the new network, and pruning non-significant connections that may have changed with the addition of a new connection (Gates & Molenaar, 2012). This process continues until the network fits the data well according to the excellent fit criteria by Brown (2015) which requires two out of four following alternative fit criteria to be met: root mean squared error of approximation (RMSEA)≤0.05, standardized root mean residual (SRMR)≤0.05, comparative fit index (CFI)≥0.95, or non-normed fit index (NNFI)≥0.95. S-GIMME identifies shared connectivity patterns while accounting for individual heterogeneity using the community detection algorithm walktrap (Beltz & Gates, 2017), which simulations demonstrate to be a reliable method of detecting subgroups of network patterns (Gates et al., 2017; Pons & Latapy, 2005) that consistently outperforms other algorithms (Gates et al., 2016). Moreover, community detection provides the model with more known priors that improves the search for individual connections (Beltz & Gates, 2017); thus, leveraging both individual and subgroup level network features increase reliability of network connections in comparison to other network approaches (Gates et al., 2017; Gates & Molenaar, 2012; Smith et al., 2011).

**Missing Data Analysis**. Prior to analysis, we assessed data missingness using the Visualization and Imputation of Missing Values ‘VIM’ package (Kowarik & Templ, 2016). We tested for Missing Completely at Random (MCAR) with the ‘MissMech’ package in R (Mortaza et al., 2014), which uses a method by Jamshidian and Jalal (2010) hat demonstrates reliability in smaller samples.

Most participants had no missing data, However, 25% of participants missing brain data (n=28). The MCAR test suggested we could not rule out MCAR (p= 0.167). Given this evidence, we concluded that estimating missing values would not introduce bias into our analysis. Using a reliable estimation method would allow us to retain power while improving confidence in our estimates. Simulations demonstrate that full-information maximum likelihood produced unbiased estimates with over 50% of missing data when missing at random (Schafer & Graham, 2002). Given that, modern missing data approaches (e.g., full-information maximum likelihood; Enders & Bandalos, 2001) reduce bias when compared to removing cases or listwise deletion (Enders, 2010; Little & Rubin, 2019), we used full information maximum likelihood to retain all 112 participants.

**Evaluating Sex as a Moderator**. Because sex is associated with CU traits and perspective taking, we assessed if sex was a moderator using multigroup models separated by sex. We constrained regression parameters across these models and compared this with an unconstrained model using a Satorra-Bentler *x^2^* difference test to determine whether model parameters are significantly different across sexes (Satorra, 2000).

**Supplementary Results**

**Zero-Order Correlations**

Zero-order correlations (Figure1) indicated perspective taking negatively associated with CU traits (r= -0.33, p< 0.01) and positively associated with cognitive control (r= 0.31, p= 0.02) and CU traits negatively associated with cognitive control (r= -0.25, p= 0.01). zero-order brain associations indicate perspective taking positively associated with positive density in the social network (r= 0.31, p= 0.01) and negative density between the social and conflict networks (r= 0.27, p= 0.03); CU traits associated negatively with negative density between the social and conflict networks (r= -0.31, p= 0.01); and cognitive control positively associated with both positive (r= 0.25, p= 0.04) and negative density in the conflict network (r= 0.26, p= 0.03; Figure 1). No zero-order associations between constructs of interest and either social or conflict network associations with the dlPFC (Figure 1).

**Sample Descriptives**

A small portion of the sample met clinical criteria for CU traits (11%; 6 male and 6 female; using criteria by Kemp et al., 2019) and conduct problems (3%; 2 male and 2 female; using criteria by Sandoval et al., 2006). Results of zero-order correlations in Figure 1 are described in Supplementary Results. In subsequent analyses, we focus on positive density in the social and conflict networks and negative density for between networks.

**S-GIMME Networks.**

Resting state networks across all participants adequately fit the data (CFI= 0.951 [min= 0.950, max= 0.956, SRMR= 0.022[min= 0.017, max= 0.028]) and were heterogenous indicated by relatively low modularity (modularity = 0.054). All person specific maps contained individual level connections (13.23±3.74) and positive connections (10.65±2.00), but negative connections were only found in 80% of the sample (2.88±1.99). All models contained contemporaneous (4.01±2.38) and lagged connections (10.32±**1.94).** Overall density suggested the social network was the most dense (social= 6.67, conflict=3.86, between social and conflict= 1.68, social and conflict with dlPFC = 1.34). A depiction of average brain network features (including centrality and connection density) can be found in Supplementary Figure 1 and for individual networks in Figure 4.

**Testing Individual Models**

***Cognitive Control Indirectly Associates with Callous-Unemotional Traits***. When building the behavioral model central to our hypothesis, we concluded that the indirect effect was central to properly representing the data, but that constraining did not impact model estimation significantly, so we removed the direct association between cognitive control and CU traits for the final model (Supplementary Table1). In this model, cognitive control positively associated with perspective taking (*β*= 0.023, *p* < 0.001, R^2^= 0.101) and perspective taking negatively associated with CU traits (*β*= -0.624, *p* < 0.001, R^2^= 0.143; Figure 2). The overall indirect effect was significant and negative (*β*= -0.014, *p* < 0.001). The model fit suggested a good fit to the data (X^2^ 8.127(8), CFI= 0.998, TLI = 0.996; RMSEA= 0.011; SRMR= 0.042; Figure 2).

***Positive Social Network Density Indirectly Associates with Callous-Unemotional Traits***. Model building determined that the indirect effects were important for model estimation but that the direct effect between positive social network density and CU traits did not impact model estimation, so it was removed (Supplementary Table 1). Model estimates indicated positive social network density positively associated with perspective taking (*β*= 2.118, *p* = 0.012, R^2^= 0.121) and perspective taking negatively associated with CU traits (*β*= -0.597, *p* < 0.001, R^2^= 0.151; Figure 2). The total indirect effect was significant and negative (*β*= -1.248, *p* = 0.021). The overall fit of the model suggested an adequate representation of the data (X^2^ 8.113(8), CFI= 1.00, TLI = 1.007; RMSEA= 0.001; SRMR= 0.058; Figure 2).

***Positive Conflict Network Density Indirectly Associates with Callous-Unemotional Traits***. Model building determined that the indirect effects were important for model estimation but removing the indirect effect between positive Cognitive Control and CU traits did not impact model estimation, so we removed this direct effect form the model (Supplementary Table 1). Model estimates indicated that positive conflict network density positively associated with Cognitive Control (*β*= 32.74, *p* < 0.001, R^2^= 0.151) and that Cognitive Control negatively associated with CU traits (*β*= -0.045, *p* < 0.001, R^2^= 0.113; Figure 2). The overall indirect effect was negative (*β*= -1.458, *p* = 0.005). The overall fit of the model suggested an adequate fit to the data (X^2^ 8.893(8), CFI= 0.974, TLI = 0.932; RMSEA= 0.046; SRMR= 0.047; Figure 2).

***Negative Between Social and Conflict Network Density Indirectly associates with Callous-Unemotional*** ***Traits***. Given we had no a priori hypothesis regarding if between network associations would indirectly effect CU traits via Cognitive Control or perspective taking, we used the results from preliminary correlations to guide our decision to focus on perspective taking as the indirect effect. Model building determined that both indirect and direct effects were important for model estimation (Supplementary Table 1). Model estimates indicated negative density between the social and conflict networks positively associated with perspective taking (*β*= 1.775, *p* = 0.005, R^2^= 0.150) and that both negative between network density and perspective taking negatively associated with CU traits (*β*= -0.464, *p* = 0.002; *β*= -2.572, *p* = 0.005, R^2^= 0.223; Figure 2). Both indirect effects (*β*= -0.703, *p* = 0.034) and total effects were negative (*β*= -3.386, *p* < 0.001). Overall model fit suggested an adequate fit to the data (X^2^ 8.291(7), CFI= 0.980, TLI = 0.939; RMSEA= 0.043; SRMR= 0.055; Figure 2).

***No Sex Differences Detected***. Multigroup comparisons stratified by sex revealed no significant difference between males and females in the study on regression parameters (Supplemental Table 4).

**Supplementary Discussion**

**No Sex Effects**

Multigroup models revealed no differences on beta coefficients across sex. This suggests that study findings may be generalized across sexes.
